# Calculation of the force field required for nucleus deformation during cell migration through constrictions

**DOI:** 10.1101/2020.12.17.423200

**Authors:** Ian D. Estabrook, Hawa Racine Thiam, Matthieu Piel, Rhoda J. Hawkins

## Abstract

During cell migration in confinement, the nucleus has to deform for a cell to pass through small constrictions. Such nuclear deformations require significant forces. A direct experimental measure of the deformation force field is extremely challenging. However, experimental images of nuclear shape are relatively easy to obtain. Therefore, here we present a method to calculate predictions of the deformation force field based purely on analysis of experimental images of nuclei before and after deformation. Such an inverse calculation is technically non-trivial and relies on a mechanical model for the nucleus. Here we compare two simple continuum elastic models of a cell nucleus undergoing deformation. In the first, we treat the nucleus as a homogeneous elastic solid and, in the second, as an elastic shell. For each of these models we calculate the force field required to produce the deformation given by experimental images of nuclei in dendritic cells migrating in microchannels with constrictions of controlled dimensions [1]. These microfabricated channels provide a simplified confined environment mimicking that experienced by cells in tissues. We extract the nuclear shape from the boundary of the fluorescently stained region in each consecutive image over time. From this we calculate the deformation field between images and use our elastic models to calculate the traction force field. Our calculations therefore predict the forces felt by a deforming nucleus as a migrating cell encounters a constriction. Since a direct experimental measure of the deformation force field is very challenging and has not yet been achieved, our numerical approaches can make important predictions motivating further experiments, even though all the parameters are not yet available. In addition, the algorithm we have developed could be adapted to analyse experimental images of deformation in other situations.

**Author summary:** Many cell types are able to migrate and squeeze through constrictions that are narrower than the cell’s resting radius. For example, both immune cells and metastatic cancer cells change their shape to migrate through small holes in the complex tissue media they move in. During migration the cell nucleus is more difficult to deform than the cell cytoplasm and therefore significant forces are required for a cell to pass through spaces that are smaller than the resting size of the nucleus. Experimental measurements of these forces are extremely challenging but experimental images of nuclear deformation are regularly obtained in many labs. Therefore we present a computational method to analyse experimental images of nuclear deformation to deduce the forces required to produce such deformations. A mechanical model of the nucleus is necessary for this analysis and here we present two different models. The first treats the nucleus as a homogeneous elastic solid and the second treats the nucleus as an elastic shell. Our computational tool enables us to obtain detailed information about forces causing deformation from microscopy images.

## 1 Introduction

Cell migration is crucial to the function of numerous cell types, for example in tissue development [2], wound healing [3, 4], immune response [5] and cancer metastasis [6, 7]. The obvious importance of cell motility in biology and medicine has motivated a large body of research. In the past, cell migration experiments tended to focus on cells crawling on rigid substrates. However in recent years, an increasing number of investigations are being done on cells in environments that are more relevant to *in vivo* tissues [8]. One aspect of tissue-like environments is that cells are confined. Studies of cells in confinement have shown that cells use different mechanisms to move than used on rigid two dimensional surfaces [9–12]. Here we concern ourselves with the case of cells migrating through micrometer sized constrictions that require significant cellular deformation.

Eukaryotic cells consist of two main compartments; the cell nucleus containing the DNA, and the cytoplasm containing the cytoskeleton, cytosol and the remaining organelles. The latter are typically smaller than the constrictions we use here and should not constitute a limitation to cell migration in this context. On the long timescales associated with cell migration, the cytoplasm displays a fluid like response to applied forces. In contrast, the cell nucleus is typically the biggest and stiffest organelle [13] and has a more elastic response [14]. The nucleus is surrounded by the inner and outer nuclear membranes, collectively known as the nuclear envelope [3]. Just inside the inner membrane is a dense intermediate filament network called the nuclear lamina [3], thought to provide the mechanical support for the nucleus and protection for the DNA [15, 16]. Recent work on cell movement through constrictions has highlighted the relevance of the cell nucleus [15–19]. The deformability of the nucleus can be thought of as a rate limiting step in cell migration through constrictions smaller than the size of the undeformed nucleus [20, 21]. In order to pass through such constrictions, a cell must generate sufficient forces to deform the nucleus such that it fits through the constriction. What force magnitudes and directions are necessary to achieve this? Measuring such forces directly is extremely challenging and has not yet been performed experimentally. However, imaging the nucleus shape is now done routinely in many labs, for example by optical imaging of fluorescently labelled nuclei. Here we therefore present a method to analyse such images to obtain predictions of the force field required to deform the nucleus in the way observed. If the mechanical properties of an object are known, the deformation caused by a known applied force can be calculated directly. However the inverse calculation to find the unknown force field that causes a known deformation is much more difficult. In fact, if only the shape outlines are known, the deformation field and forces cannot be calculated exactly. This is because there is no unique mathematical mapping between two shapes if only the outlines are known. In this work we therefore develop a numerical simulation method to calculate the force field based on physical modelling.

In this study we consider the case of cells migrating through microfabricated channels [1] (see Fig 1). In this setup there are no external flows or chemotactic gradients and the spontaneously motile cells move themselves through the channels, entirely self-generating the required forces. The channels are made of polydimethylsiloxane (PDMS) with a glass bottom through which the microscopy images are taken. All surfaces of the enclosed channels are coated with fibronectin. We choose the height and width of the microchannels to match the cell size in order to effectively constrain cells to one dimensional motion along the long axis of the channel. We design constrictions in the channels such that cells and their nuclei must deform to pass through the constrictions. We obtain sequential images of dendritic cells with DNA fluorescently stained with Hoechst as the cells migrate through constrictions in the microchannels. From these images we extract the shape of the nucleus at each time point.

**Fig 1.**
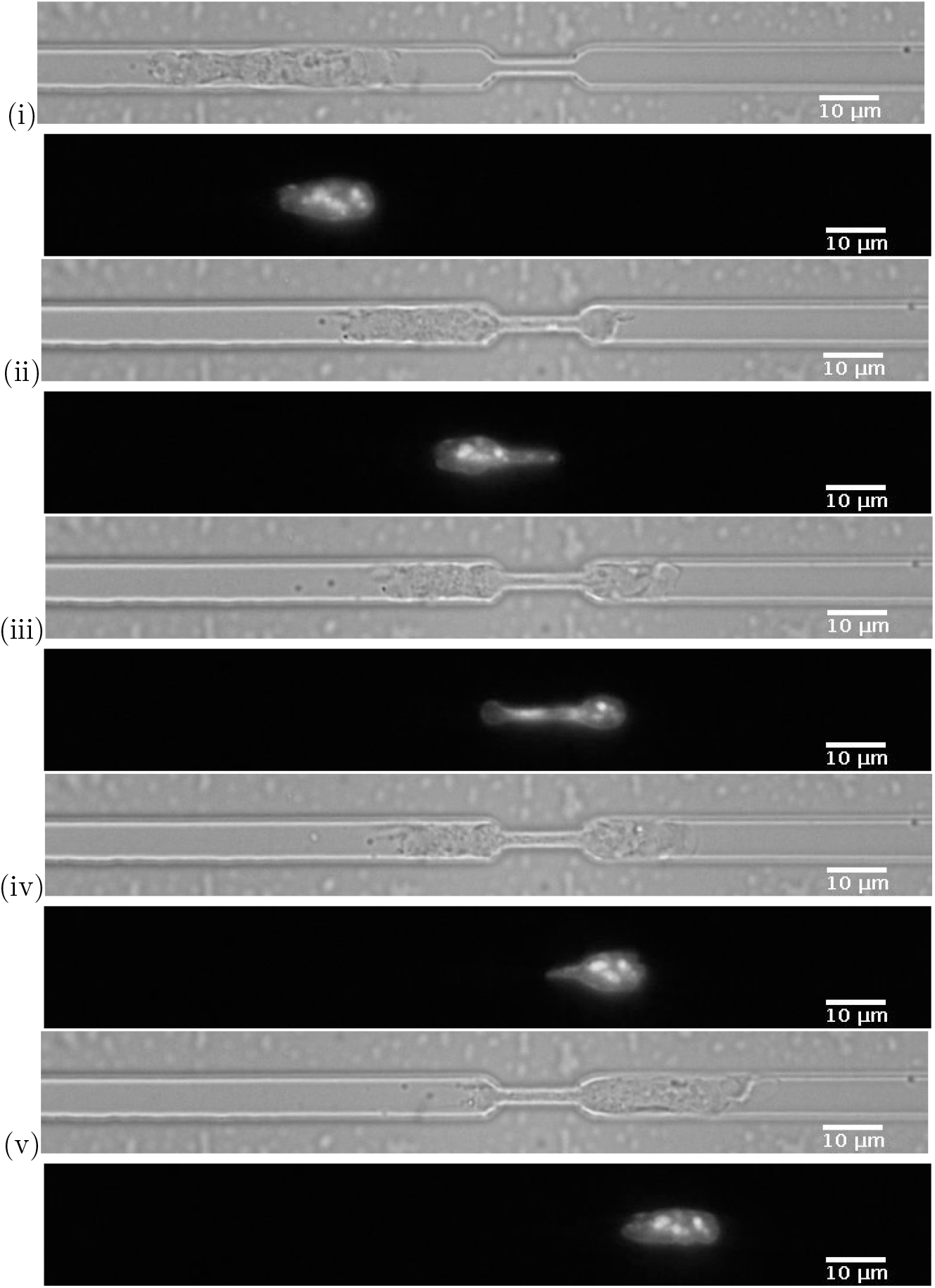
(Top) Confocal microscopy image of an example mouse dendritic bone marrow cell, passing through a constriction in a microfabricated channel at different time points (i)-(v). (Bottom) Fluorescent microscopy image of the nucleus of the same cell, stained with DAPI. The scalebar is 10 *μ*m and the constriction width in this image is 2 *μ*m. The chosen time points are; (i) before the constriction, (ii) beginning to enter the constriction, (iii) within the constriction, (iv) exiting the constriction and (v) after the nucleus has fully exited the constriction.

A mechanical model is necessary to deduce the force fields causing the nucleus deformations observed in experiments such as the microchannel experiments we focus on. To find a deformation field between the different nucleus shapes obtained from the experimental images, we use a mechanical model and a simulated annealing approach. We then use our mechanical model to calculate the force field required to produce the deformation. This gives us a prediction of the forces generated by the cell to deform the nucleus through constrictions. These predictions are dependent on the mechanical model for the nucleus used. In this work we consider two simple limiting cases of possible models for the nucleus. However, the algorithm we have developed can be used with any mechanical model and therefore is easily extendable to more realistic models for the nucleus.

To model the nucleus we assume it is an elastic material, governed by the laws of continuum elasticity. We consider two different single component models for the nucleus, namely a homogeneous elastic solid and a thin elastic shell, which treat the extremes of inner material behaviour (see section 2).

To date a few groups have published models for the nucleus using a variety of approaches. The simplest is a quasi one dimensional model of a linear elastomer [22]. More complex simulation models include a liquid droplet with an elastic interface using the immersed boundary method [23], cellular Potts with an energy cost to deformation [24], finite element analysis of a viscoelastic [25] or poroelastic [26] material and hybrid agent based finite element [27]. The approach closest to ours in spirit is that of [28] who use an elastic model to analytically calculate the energy required to deform the nucleus from an initial spherical shape to an ellipse or a cigar shape. A common feature of all these previous models is that the deformation is imposed by the model, whereas our model calculates the deformation from inputted arbitrary shapes extracted from experimental data. Due to the complexity of experimentally derived shapes we have to calculate the deformation field using numerical simulation. Results from these previous studies indicate under what circumstances a cell is able to enter a constriction. Our model however calculates the force field required for the deformation observed experimentally. In other words, the force exerted on the nucleus is the output of our model not an input.

The rest of the article proceeds as follows. We discuss the physical properties of the nucleus in more detail in section 2. Then in section 3 we introduce the continuum elasticity formalism and present our homogeneous elastic solid and elastic shell models. We describe in section 4 the Monte Carlo procedure we use to determine the deformation field from the experimental images. We discuss our results in section 5 and finally conclude with section 6.

## 2 Physical properties of the cell nucleus

In this section we discuss the mechanical properties of the nucleus focusing first on the elasticity of the nucleus and then on its compressibility.

### 2.1 Nucleus elasticity

In general the elasticity of the nucleus might depend on the timescale and on the amplitude of deformation. The nucleus appears elastic on the minutes timescales we are concerned with, as evidenced by Neelam et al. [29] who showed that the nucleus regains its original shape within seconds after removing a deforming force. In the microchannel experiments we analyse, following exit from the constriction the nucleus regains its original shape immediately (within the time resolution of our experiments), implying it is behaving elastically. Returning to the original shape might in general involve active mechanisms such as ion pumps, however such processes are slow and we assume slower than the timescale of the nucleus response seen in our system [30]. The cell nucleus has been seen to display a significantly more elastic response to applied forces than the surrounding cytoplasm in micropipette aspiration experiments on a timescale of minutes [14, 31]. The mechanical properties of the nucleus have also been investigated using atomic force microscopy, in which indentation caused by small applied forces demonstrate that the nucleus is stiffer than surrounding cytoplasm [32]. The nucleus tends to be an order of magnitude stiffer than the rest of the cell in healthy cells [18]. The elastic response of the nucleus is thought to be provided mainly by the nuclear lamina, in particular by lamins A and C [29, 33–36]. The lamina is a ~ 100 nm thin network of intermediate filaments (lamins) on the inner side of the nuclear envelope [3].

The response of the internal elements of the nucleus is not so clear. Stem cells not expressing lamin A/C show plastic deformation [37]. Nuclei lacking lamins appear viscoelastic if chromatin is tethered to the nuclear membrane but are viscous-like if chromatin is untethered [38]. Recent micromanipulation experiments on isolated nuclei suggest that chromatin dominates the mechanical response to small deformations but the lamin A/C shell dominates the response at large deformations such as those experienced during migration [39]. Our two models treat the extremes of inner material behaviour (elastic or completely deformable). In reality we expect the nucleus inner material may be viscoelastic and thus have properties between the two extremes we address in this work.

We characterise the nuclear elasticity by the Young’s modulus and Poisson ratio (see section 2.2 for discussion of the latter). We choose a value of 5 kPa for the Young’s modulus of all our nuclei, which is consistent with the reported range of values in the literature [31, 40, 41]. However, since the stress and traction force field are proportional to the Young’s modulus (see Eqs in section 3), our results can be easily scaled for different values of the Young’s modulus. The magnitude of the forces is directly proportional to value of the Young’s modulus but the directions of the forces are independent from the value of the Young’s modulus.

### 2.2 Nuclear compressibility

To model the nucleus we need to know its compressibility as well as its elasticity. Models often assume incompressibility but some experimental studies have indicated that the nucleus can be compressible [30, 42, 43]. However whether the nucleus is compressible may depend on timescale, amount of deformation and cell type. Therefore we investigate whether the nucleus undergoes significant volume changes during deformation in our experimental situation. We estimate the volume by measuring the area of the central plane from the experimental images and making assumptions about the geometry in the third dimension. We measure the area (in the plane of the microscope, *x* − *y*) of the nucleus of each cell as it migrates through the constriction (data shown in Appendix 6.1 Fig 7). The nucleus area increases as it goes through the constriction before returning to its pre-entry area, which itself undergoes small fluctuations. To determine the volume we require information about the out of plane direction, *z*. Fig 2(b) shows a *z*-stack of images of *x* − *y* cross sections taken at different heights, *z*. The observed constant cross section indicates that the nucleus fills the channel and therefore also the constriction. We therefore calculate an estimate of each nucleus volume from the known channel geometry. The channel cross section dimensions (width × height) are 7*μ*m × 4.7*μ*m outside the constriction, and 2*μ*m × 3.4*μ*m inside the constriction. We estimate the nucleus volume by calculating the area within and without the constriction and multiplying it by the respective heights. Fig 2(a) shows the mean volumes of each nucleus before, inside and after the constriction, showing there is no clear volume increase or decrease as nuclei pass through the constriction. There is no statistical difference between the mean volumes over all the nuclei before the constriction (240 ± 90) *μ*m^3^, in the constriction, (223 ± 67) *μ*m^3^ or after the constriction (235 ± 57) *μ*m^3^. The change in projected density of DNA, as determined by the fluorescence intensity, is also consistent with the channel height change in the constriction (see Appendix 6.1).

**Fig 2.**
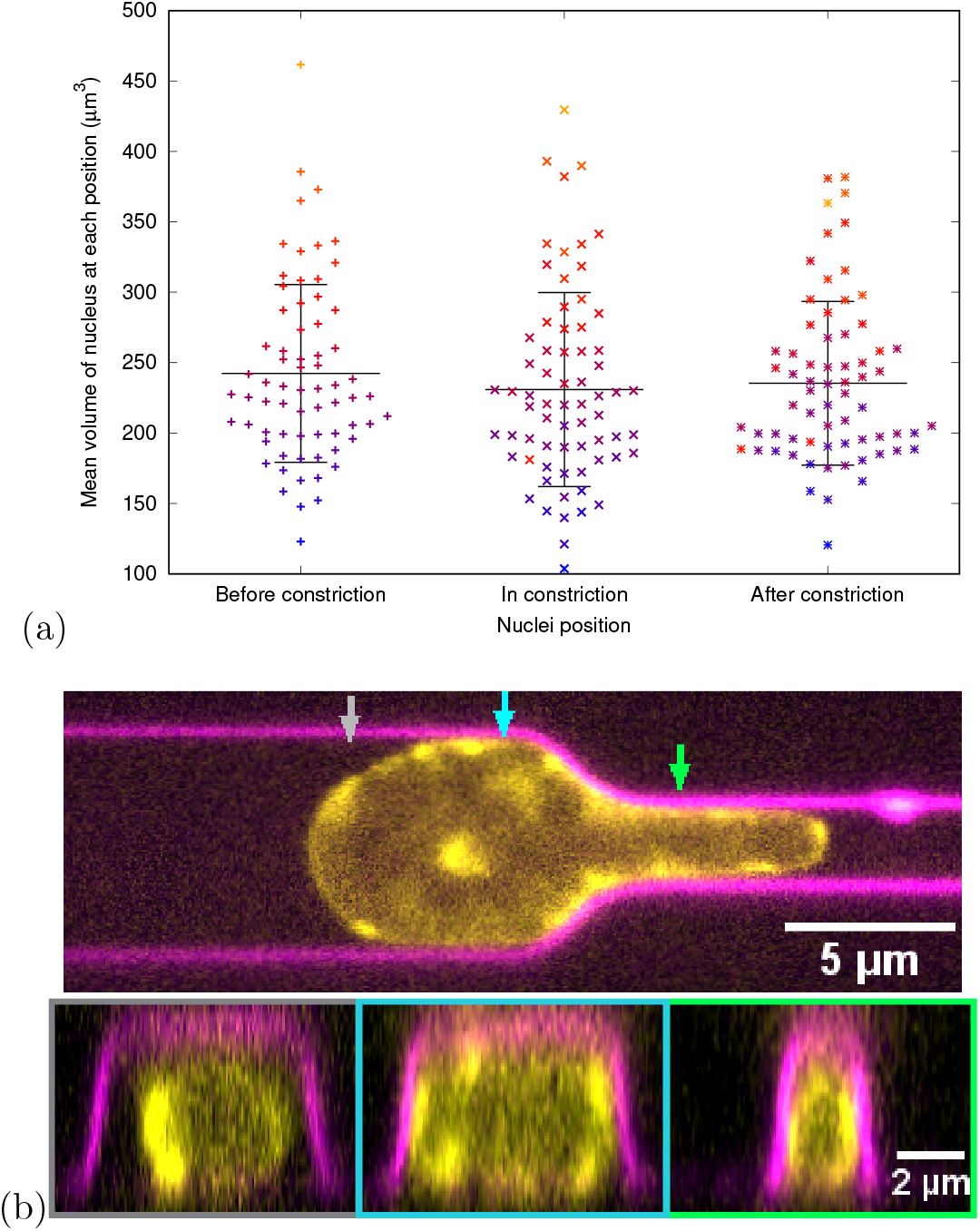
(a) The mean volume of each nucleus in *μ*m^3^ before the nucleus enters the constriction, while any part of the nucleus is between the constriction entry and exit, and after the nucleus has exited the constriction. Each mean value is calculated using every time frame at the corresponding position of the nucleus relative to the constriction. Each point is an individual nucleus with an identifying colour. The colour scale blue to red to orange matches the volume of the nucleus before the constriction from small to large. Nuclei in the later positions keep their colour as defined by their volume before the constriction. The fact that the colours of the points stay mostly in the same order shows that the volume of a particular nucleus is roughly constant. The black horizontal lines show the mean volume of nuclei at that position and the vertical black lines indicate the standard deviations. Clearly there is no significant difference in mean nuclear volume as a nucleus travels through the constriction. (b) Side view nucleus in a channel. Spinning disc confocal image of a dendritic cell with the DNA (in yellow) stained with Hoechst, migrating through a channel with a 2*μ*m wide and 20*μ*m long constriction. Channels are coated with mCherry-PEG (magenta). Up image: top view of the channel. Bottom image: orthogonal/side view of the channel. Arrows (grey, cyan and green) in upper image to indicate the slicing regions depicted in the bottom image surrounded by grey, cyan and green rectangles.

From these measurements we conclude that the nuclei in our experiments do not significantly change volume during deformation through the constrictions. In our models we therefore treat the nucleus as an incompressible material with Poisson ratio *v* = 0.5.

## 3 Continuum elasticity

We treat the nucleus as an elastic object, as assumed for example by [28, 40, 44] (see also discussion in section 2.1). We use a continuum approach, justified by our interest in the properties of the nucleus as a whole on length scales much larger than the constituent molecules.

The standard constitutive equation for a continuum linear elastic solid obeying Hooke’s law is given by [45]

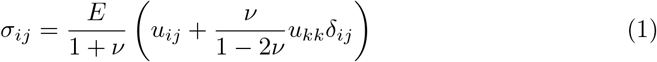

where *σ*_*ij*_ is the stress tensor, *E* is the Young’s modulus, *v* is the Poisson ratio and *u*_*ij*_ is the strain tensor. The subscripts label tensor elements and *δ*_*ij*_ is the Kronecker delta function. We use the Einstein summation convention for repeated indices. The strain tensor is defined by:

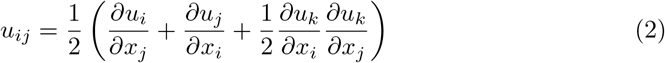

where *u*_*i*_ is the displacement and *x*_*i*_ is the position vector. Note that here we keep the second order terms, which are often neglected for small deformations. The free energy density, *f*, of a deformed elastic solid expressed in terms of the strain and stress tensors is given by [45];

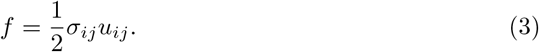

The traction force is defined as the external force on a unit area of the surface of a body. For an elastic object it is given by [45];

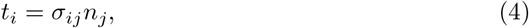

where *n*_*i*_ are the components of the normal to the surface.

In the following we present two elastic models; treating the nucleus as a homogeneous elastic solid (section 3.1) and as a thin shell (section 3.2).

### 3.1 Homogeneous elastic solid model

We first consider the simplest model of a nucleus as a single homogeneous elastic solid. This assumes the inner nuclear material is elastic on migration timescales and can be thus treated in the same way as the lamina. The is the fully elastic extreme of the models we consider and is useful for comparison to the shell model and to provide the upper limit of the forces required for deformation. Since the strains in our case are not sufficiently small to neglect the nonlinear term we take the full expression given in Eq 2.

To treat the nucleus as an elastic solid, we require the deformation field within the nucleus. Due to the limits in spatial and temporal resolution and lack of clear landmarks, we cannot obtain this directly from the experimental images. We therefore extrapolate the deformation field inside the nucleus from the deformation of the experimentally obtained nuclear outlines. We consider deformations in the centre of mass frame of the nucleus. We assume the centre of mass has zero deformation between any two given images, i.e. the deformation field at (0, 0, 0) is (*u*_*x*_, *u*_*y*_, *u*_*z*_) = (0, 0, 0). We then assume that the deformation field decreases linearly along the radii from each boundary point to zero at the centre of mass. We calculate the centre of mass of a nucleus from its input outline by using the approximation of the centre of mass of a general polygon, given by [46],

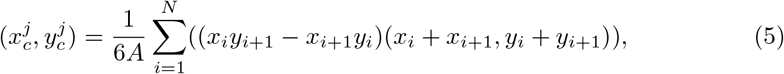

where 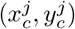 are the coordinates of the centre of mass of frame *j* of the video, *A* is the area of the polygon, and (*x*_*i*_, *y*_*i*_) are the coordinates of each of the *N* points making up the polygon representing the outline of the nucleus. The polygon is a closed loop such that the final point *N* connects directly to the initial point *i* = 1.

To numerically calculate derivatives in polar coordinates we use central/forward/backward finite difference methods for the middle/innermost/outermost rings respectively (see Appendix 6.2 for details). Along with the standard relations between Cartesian and polar coordinates, this allows us to evaluate the strains directly in the Cartesian coordinate basis using Eq (2).

Fig 2(b) shows that the cell nucleus contained in a channel has the same cross section at different heights, *z*. Therefore we assume that the nucleus curvature in the z direction is zero, both within the channel and inside the constriction. Around the entrance and exit of the constriction where the height is changing there will be a non-zero curvature in the z direction. However, this does not affect our calculations since we calculate traction forces in the *x* − *y* plane which are independent from those in the *z* direction. We assume that the images of cell nuclei in the *x* − *y* plane (such as figure 1) correspond to the central plane of the nucleus, *z* = 0. We use the incompressibility condition to determine the strain component in the *z* direction, *u*_*zz*_ = −(*u*_*xx*_ + *u*_*yy*_), where the strain components *u*_*xx*_ and *u*_*yy*_ are calculated from the deformation fields of the images in the *x* − *y* plane. Since the deformation in the *z* direction, *u*_*z*_, varies only along the channel direction, *x*, the strain components *u*_*yz*_ = *u*_*zy*_ = 0. In the central plane, at *z* = 0, the deformation *u*_*z*_ = 0 and strain components *u*_*xz*_ = *u*_*zx*_ = 0.

Whilst in general a non zero *σ_zz_* stress term exists, the curved surface of the nucleus is defined so that its normal is perpendicular to the 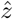 direction. Therefore the *σ*_*zz*_ term does not contribute to the traction force on the surface seen in images. The traction force, Eq (4), on the central plane can then be solved based on the two dimensional problem, since the traction over the central plane is unaffected by the surfaces at *z* ≠ 0.

### 3.2 Thin elastic shell model

In this section we present our model of the cell nucleus as a thin elastic shell. We briefly describe the differential geometry needed to calculate the strains here and give further details in Appendix 7. We treat the nucleus as a thin elastic shell assuming the inner part of the nucleus contributes no resistance to deformation. This means that in principle the inner medium could change volume, however in our case the observed deformations do not significantly change volume (see section 2.2). Our shell model is motivated by the different responses to deformation of different components of the nucleus, as described in section 2. Specifically, we treat only the nuclear lamina as having an elastic response to deformation and assume the inner nuclear material provides no resistance to deformation. Therefore we do not need to determine the deformation inside the body, as was required for the solid model in section 3.1.

Treating the nuclear lamina as an elastic shell is motivated by the assumption that the elastic response of the nucleus is provided mainly by the nuclear lamina, in particular by lamins A and C [29, 33–36]. We assume the nuclear lamina is a homogeneous elastic material with the same elastic parameters as we used for the whole nucleus in the solid model (section 3.1). The amount of any pre-stress in the shell or excess area prior to entry to the constriction has not been established experimentally. Therefore, for simplicity, we assume there is no pre-stress nor excess area.

The nuclear lamina is only about ~ 100 nm thick [3, 47], which is thin compared to even the smallest constriction width of 2 *μ*m. We therefore use a thin shell approximation in the calculations below. We follow the Love-Kirchhoff thin plate theory [48] as used by Berthoumieux et al. [49] for their model of the cell actomyosin cortex. This thin shell approximation assumes the thickness of the shell remains constant and the normal to the surface remains normal after deformation. This means that the only non-zero strains are those in tangential directions. It is therefore convenient to calculate the strains in the normal and tangential coordinate basis, before transforming back to Cartesian coordinates.

As drawn in figure 3, we define the normal and tangential coordinates as (*n, s*_1_, *s*_2_) where *s*_1_ is along the tangent to the surface in the plane of the images 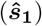, *s*_2_ is along the surface in the out of plane direction 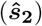 and *n* is along the outward normal to the surface 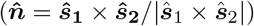. We assume the shape is symmetric in the out of plane direction and therefore that the normal, 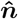, is in the *x*-*y* plane and the tangent 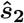 is along the 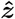 direction. This means our coordinate basis is orthonormal and therefore the metric tensor is the identity matrix and all Christoffel symbols are zero so the derivatives of the basis vectors along the surface are given by the curvature tensor, *C*_*ij*_, i.e. 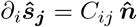, where the curvature tensor 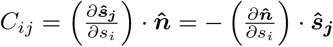. We assume that the surface of the nucleus has zero curvature in the out of plane direction (apart from at the constriction entrance and exit), as motivated by the assumption that the nucleus is symmetric and fills the channel at all times, as in section 2.2. As discussed in section 3.1 the surfaces at *z* ≠ 0 do not affect the traction forces in the *x* − *y* plane. In this case the only non zero term in the curvature is 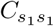 which is calculated numerically from the image data.

**Fig 3.**
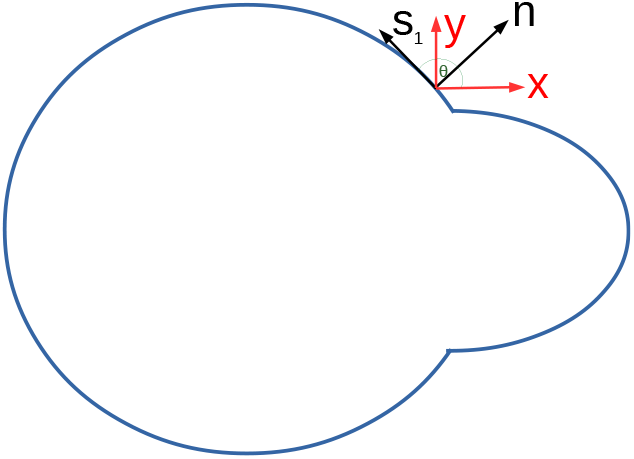
A cartoon of a deforming nucleus in 2D showing the coordinate basis (*n, s*_1_, *s*_2_) used for the shell model compared to the Cartesian coordinate system used in the 2D images.

The deformations of surface points are initially calculated in Cartesian coordinates. We transform these to our normal and tangential basis using standard clockwise rotation matrices *R*_*θ*_ as ***u***(*n, s*_1_, *s*_2_) = *R*_*θ*_*R*_*ϕ*_***u***(*x, y, z*) where *θ* is the angle between 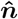 and 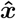 and *ϕ* is the angle between 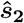 and 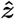, which in this case is *ϕ* = 0. Using this basis, we can calculate the strain, Eq 2, and then transform back to Cartesian coordinates; 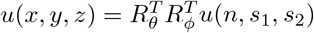. We then calculate the stress and traction from Eqs (1) and (4) as for the solid model.

Note that, as for the solid model, we assume the traction forces on the central plane are unaffected by the edges at finite *z* values and therefore we do not consider the part of the shape with components to the the normal in the 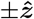 direction.

## 4 Calculation of the deformation field

We calculate the deformation field in the central *x*-*y* plane from the 2D experimental images at consecutive time points. We first find a list of coordinates defining the outline of each image using the image analysis software ImageJ [50]. We convert the image to binary using the set threshold tool and then use the analyse particles tool to obtain the nuclear outline curves at each time point. We choose the threshold to distinguish between the class of bright pixels that are in the nucleus and those that are in the noise far from the nucleus. We do this by plotting the histogram of number of pixels against intensity and setting the threshold value to minimise the intra-class variance (Otsu’s method). Finally, we use the spline fitting and interpolation tools to obtain a list of boundary coordinates, spaced one pixel apart. The list of points on this curve is then used as the input to our algorithm described below.

### 4.1 Averaging of different images

To reduce the effects of noise in the data, we averaged data for many cells (between 55 and 71 with sufficient resolution). To save computational time we averaged the input shapes and then calculated the forces required for these averaged nucleus deformations. We averaged the nuclei according to position relative to the entrance of the constriction. The points we chose are; the nucleus prior to entry (the frame before the nucleus begins deforming in to the constriction), the nucleus entering (chosen as the first frame where the nucleus begins deforming into the channel), the nucleus entirely within the channel (the first frame where the rear of the nucleus completely passes the constriction entrance), the nucleus exiting (the first frame where the front of the nucleus passes the constriction exit) and the nucleus after leaving the constriction (the first frame where the rear of the nucleus has moved past the constriction exit). The average outline is obtained by first finding the geometrical centre of each individual outline and aligning the images with their centre of mass at the origin. Note that in our model the centre of mass is at the same point as the geometrical centre. We then draw spokes radially outwards from the centre of mass to the outline at fixed angular values, every 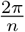, where *n* is the average number of pixels in the outlines. Then the coordinates of each spoke at the same angle for each shape are summed up and divided by the number of input shapes to give the final average nucleus shape for that time point.

### 4.2 Alignment of nuclear images to remove translation

To define the deformation field between two images of cell nuclei under motion, we need to choose a point to register the images. Our aim is to remove the translational motion of the nuclei, so that the forces we calculate are only those causing the shape change of the nuclei. Therefore for the solid model we set the centre of mass of the initial shape to have zero deformation and we assume deformations at the surface of the nuclei linearly drop to zero at the centre of mass. However, unless the deformations are symmetric, the centre of mass of the target deformed shape will not be at the same point as the centre of mass of the initial shape. Therefore, we cannot, *a priori* assume the images should be aligned with the nuclei centre of masses coinciding. When the nucleus starts to enter the constriction the rear moves forwards very little compared to the leading edge. However, we cannot *a priori* assume the images should be aligned with the rear of the nuclei coincident.

The correct alignment point is likely to lie somewhere between the two limiting cases of the centre of masses or the rear/front points. The limits on both the spatial and temporal resolution of the nuclei in the images prevent us from directly identifying the point of zero deformation from a given video. Therefore, to estimate the translation free relative alignment of the nuclei, we measure the change in the front and rear positions of each nuclei between time frames. We set the centre of mass of the initial shape to be at the origin. We then shift the target shape along the *x* axis a distance proportional to the measured relative change in the rear and front positions. See Appendix 8 for details.

### 4.3 Simulated annealing

In this section we describe the Monte Carlo simulated annealing method we use to determine the deformation field between two images of a cell nucleus. The algorithm we have developed reads in a list of coordinates describing the perimeter line of a 2D image for each time step. In our case these coordinates are obtained from experimental images of nuclei, an example of which is given in Fig 1(b). However, the same algorithm could be used for other elastically deforming objects. Our aim is to determine the deformation field between two images of the cell nucleus boundary. However, mathematically there is no unique mapping between two such images. We assume that the mapping describing the physical deformation is the one that minimises the free energy of deformation, given by Eq (3). To calculate this energy minimised deformation field, we use a simulated annealing approach, as described below.

In order to determine the deformation field between two images of cell nuclei, the nucleus must be tracked from image to image. One common method to determining the deformation fields between images is by following known ‘landmarks’ in the images [51, 52]. A landmark is a recognisable area that can be used to determine a local deformation field. The global deformation field is then extrapolated from the local deformation of landmarks. However, due to the complex nature of the nucleus and the limited resolution currently available, there are no consistently reliably identifiable landmarks in the Hoechst stained images of the nucleus described above. A different approach is needed in order to determine the deformation field from images lacking clear landmarks. We therefore determine the deformation field through an energy minimisation simulated annealing routine. That is, we seek the mapping that minimises the free energy of the deformation, given by Eq (3), and we assume that this mapping describes the physical deformation the nuclei undergoes. We calculate the deformation field for the outline of the nucleus from each image, as described below.

Consider two input image boundaries at consecutive time points. The nucleus deforms from the first image shape to the second. We assume that boundary points maintain sequential order during any deformation. The first image we will refer to as the initial image and the second as the target image. We first choose an arbitrary mapping between the initial and target image boundaries to obtain an approximation to the deformation field. We will then refine this using a simulated annealing approach to find the deformation field that minimises the free energy. To define the arbitrary mapping, we first order the list of boundary pixel coordinates for each image by angular position and then remesh the initial boundary so it has the same number of points as the target boundary, at equally spaced angles. We calculate the free energy of this arbitrary starting deformation using Eq (3).

We now perturb the deformation field to find configurations with lower free energy. We randomly choose one of the initial pixels and perturb the position of its mapped point on the target image. We move the chosen pixel a tenth of the way towards one of the two neighbours, with the direction chosen at random. We calculate the free energy of this new configuration. If the free energy of deformation decreases, the new deformation location is recorded as a new minimal energy configuration. If the free energy increases the energy by an amount Δ*E*, the configuration is kept if 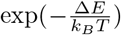 is greater than a randomly generated number between 0 and 1. If not, then the change is discarded, and a new perturbation performed. Perturbations are continued until a minimum energy is reached. The temperature *T* is gradually reduced during the simulated annealing procedure from an initial value such that 20% of moves are accepted, as in standard simulated annealing procedures, which are described in more detail in [53, 54]. In principle this procedure could fail if there was more than one energy minimum. However, the procedure works well for small deformations within the validity of the elasticity model we use. Our method would fail for deformations beyond the elastic limit (for example a ruptured shell) or for deformations causing the surface to bend back upon itself causing multi-valued surface points at a particular angle. We find it works well for the types of deformation we study here.

## 5 Results and Discussion

The nucleus is now considered to be the main element hindering cell migration through tissues [18]. It is therefore crucial to understand how cells deform the nucleus, especially professionally migrating cells such as dendritic cells. In this section we present our results for the traction force field on the nucleus calculated from the deformation field of average shape changes calculated as in section 4.1, using the aligning estimated as described in section 4.2 and the simulated annealing method described in section 4.3. We present and compare results for both our homogeneous elastic solid (section 3.1) and thin elastic shell (section 3.2) models.

### 5.1 Solid model results

Fig 4 shows the solid model deformations and traction force fields between the average nuclear shapes at the time points chosen with respect to the constriction; before to entering (a), entering to fully inside (b), fully in to exiting (c) and exiting to fully after (d). Note that in our elastic model, the Young’s modulus, *E*, linearly scales the magnitudes of the traction forces without affecting their directions nor distributions. In figure 4(a), the nucleus begins to enter the constriction. The resulting traction force shows compression everywhere except close to the rear and front of the nucleus where the forces are outwards. This is to be expected since the width of the nucleus needs to narrow to enter the constriction. The largest magnitude forces (~ 4 kPa) are at the leading edge. The traction force as the nucleus goes from entering to fully inside the constriction, shown in figure 4(b) has a positive *x* component almost everywhere. This is predictable, given that the front of the nucleus moves more compared to the rear of the nucleus during this step than in any of the other time steps (see figure 8b). Fig 4(c) shows the traction force as the nucleus changes shape from inside the constriction to beginning to exit. Apart from the rear and the very front, the traction forces are outwards as the shape widens on exiting the channel. These outward traction forces are larger in magnitude near the front than the rear, due to the larger deformation of the front than the rear as the nucleus exits the constriction. Also noticeable particularly in figure 4(c) is that the energy minimisation process results in points making up the surface being closer together in the centre than at the ends. This is due to the fact that the centre of mass of the initial shape has zero deformation. The final figure panel, 4(d) shows the traction force on the nucleus as it changes shape from the average during exit of the constriction, to the average when the nucleus has fully left the constriction. The average shapes used are notably similar to the shapes taken before the constriction and on entering, as we would expect from an object displaying elastic behaviour. The magnitude of the force is now significantly larger at the rear than at the front, which coincides with the rear of the nucleus now deforming more to regain the shape prior to the constriction, whilst the front of the nucleus has already left the constriction so deforms less at this point.

**Fig 4.**
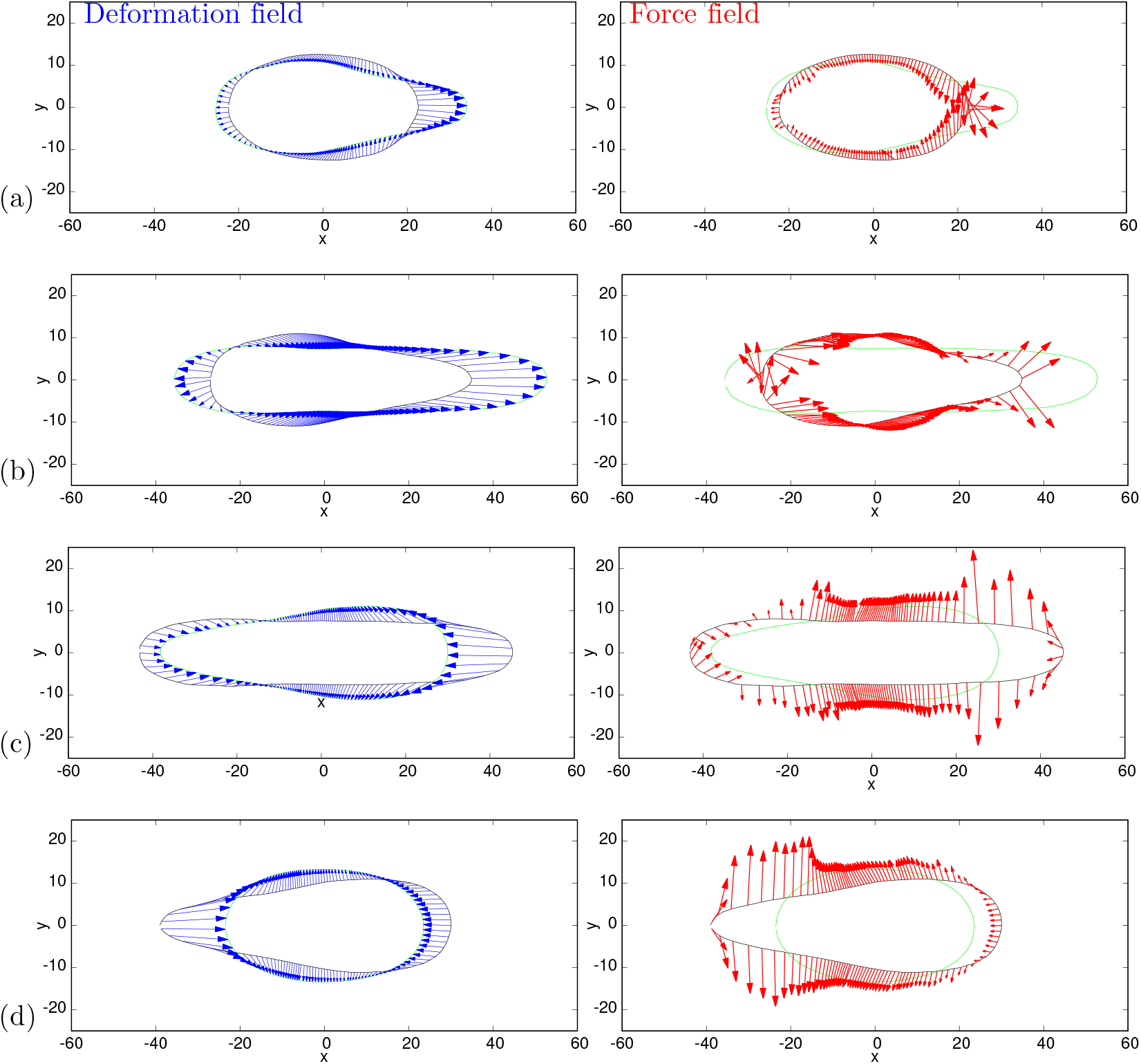
Deformation (left) and traction force (right) fields for the solid model. The axes show pixel numbers where each pixel is 0.215*μ*m. The black outline is the initial shape and the green outline is the target deformed shape. Blue arrows represent the final deformation field found between the images and red arrows represent the traction force direction and magnitude, with each arrow scaled such that one unit of length on the axes represents a traction force of 250 Pa. The traction force is calculated using a Young’s modulus of *E* = 5 kPa and assuming that the nucleus behaves as an incompressible elastic solid with a Poisson ratio *v* = 0.5. (a) Initial shape is the average of 71 nuclei before the constriction and the target shape is the average of 56 nuclei beginning to enter the constriction. (b) Initial shape is the average of 56 nuclei entering the constriction and the target shape is the average of 71 nuclei within the constriction. (c) Initial shape is the average of 71 nuclei within the constriction and the target shape is the average of 55 nuclei as they begin exiting the constriction. (d) Initial shape is the average of 55 nuclei exiting the constriction, to the average shape of 71 nuclei after they have fully exited the constriction. The target shape is moved +1.7, +7.3, +0.15, 0.0 pixels along the *x* direction, from the centre of mass aligned position for (a)-(d) respectively (see Appendix 8).

Large compressive lateral inward forces with components perpendicular to the channel axis are clearly seen in Fig 4(a) and (b) exerted on the nucleus when it enters the constriction. This is quite different from the axial force expected from the common view that cells push the nucleus through the constriction by actomyosin contractility at the back building up a pressure difference between the back and front [8, 28, 55]. The competing idea that the cell pulls the nucleus from the front (using either actomyosin contraction and connection to the substrate at the front, or microtubules connecting the nucleus to the cell front) would also generate axial forces. If we consider the cell generates a force parallel to the channel axis as it does in both push and pull models and in cell migration, the normal reaction force of the sloped walls at the entrance to the constriction will have a component inwards perpendicular to the channel axis. However, the inward force magnitudes calculated are larger than that expected for the component due to the passive reaction of the sloped walls to a forward migratory force. In Fig 4(a) some of these lateral forces are even directed backwards and therefore cannot be only passive reaction forces balancing a forward cell generated force. From Fig 4(b) and (c) it is clear that inward compressive forces on the nucleus also occur inside the constriction perpendicular to the walls and channel axis. These forces cannot be due to a reaction force to a cell generated force parallel to the axis of motion. This therefore implies an additional active force generated by the cell to compress the nucleus on entry to and in the constriction. Thiam et al. [17] proposed that an additional force might be important, with actin polymerisation compressing the sides of the nucleus. Whilst Thiam et al. [17] gave clear evidence that this actin was important for nuclear passage, they provided no direct proof, nor an explanation why it would be important. Here we show directly that active forces are deforming the nucleus from the sides. In section 5.3 we compare our results with the actin measured by Thiam et al. [17] providing the first evidence that this lateral actin is exerting forces deforming the nucleus.

The outward forces in Fig 4(c) and (d) are indicative of an elastic response of the nucleus favouring a return to its original shape once the nucleus leaves the constriction. These outward forces perpendicular to the channel walls seen in Fig 4(c) and (d) correspond well with the forces implied by the reflection interference contrast microscopy (RICM) measurement by Thiam et al. [17]. They found by RICM that the only parts the cells in close contact with the channel walls are the cell rear and inside the constriction. Fig 4(c) and (d) suggest that close contact with the channel walls would be seen when the nucleus is in, and exiting from, the constriction.

### 5.2 Shell model results

Fig 5 shows the deformation and traction force fields calculated using the shell model for the same time points and average nuclear shapes as shown in figure 4. The deformation fields are slightly different from those in figure 4 due to the model dependence of the deformation energy and therefore the minimisation procedure. In the solid model (figure 4), points on the target shape are positioned to minimise radial distance whereas in the shell model (figure 5), the deformation minimises the number of points in regions of high curvature. This is due to the fact that in the shell model the strain is dependant on the curvature due to the derivatives of the basis vectors (see section 3.2). Therefore, in the high curvature regions the energy minimisation results in larger deformations between neighbouring points, the cost of which is outweighed by minimising the energy cost due to curvature. This physical effect means the high curvature regions are more stretched and therefore more likely to rupture. We have not included the possibility of rupture in our model but one could consider rupture occurring above a threshold tension.

**Fig 5.**
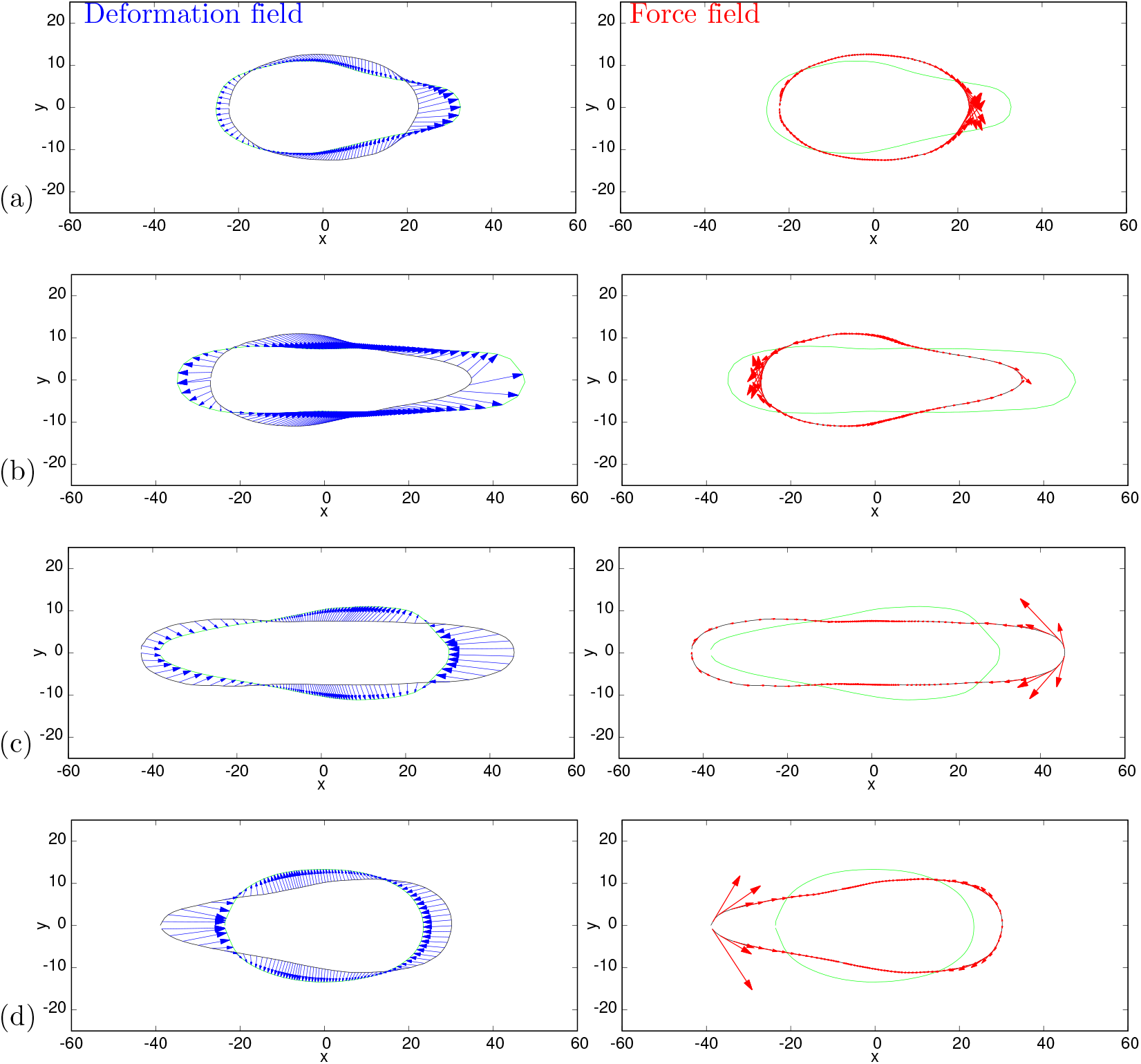
Deformation (left) and traction force (right) fields for the shell model. The axis show pixel numbers where each pixel is 0.215*μ*m. The black outline is the initial shape and the green outline is the target deformed shape. Blue arrows represent the final deformation field found between the images and red arrows represent the traction force direction and magnitude, with each arrow scaled such that one unit of length on the axes represents a traction force of 250 Pa. The traction force is calculated using *E* = 5 kPa and assuming that the nucleus behaves as an incompressible elastic shell with a Poisson ratio *v* = 0.5. Average shapes the same as in Fig 4, i.e. (a) before to entering constriction, (b) entering to within constriction, (c) within to exiting constriction and (d) exiting to after constriction. As in Fig 4, the target shape is moved +1.7, +7.3, +0.15, 0.0 pixels along the *x* direction, from the centre of mass aligned position for (a)-(d) respectively (see Appendix 8).

All the traction forces in figure 5 are parallel to the surface. This is because in the thin shell approximation the normal component of the strain is zero and, due to the incompressibility condition, a zero normal strain leads to a zero normal stress. Therefore the strain and traction force are purely tangential. In general, the magnitudes of the traction forces are smaller in the shell model (mean 0.5 kPa) than for the equivalent elastic solid model (mean 1.5 kPa) shown in figure 4. This is expected given that for the shell model the inner material has no mechanical resistance so the force required to deform a thin elastic shell is less than that required to deform a solid elastic object.

Of particular interest is the traction force in figure 5(c) which shows large extensile traction forces at the leading tip when the nucleus is at its most elongated inside and exiting the constriction. These forces are suggestive of the forces required to break the lamina [17] and induce nuclear envelope rupture as observed at the front tip of nuclei in small constrictions [15, 16]. Note we do not expect the large forces at the rear seen in figure 5(d) to also cause rupture since these forces occur at the time point when the rear of the nucleus exits from the constriction so the nucleus is then free to reform a less confined shape thus preventing rupture.

### 5.3 Correlation with actin density

The lateral inward forces shown in Fig. 4(a) and (b) are suggestive of forces generated by the Arp2/3 mediated actin assembly around the nucleus when it is in the constriction reported by Thiam et al. [17]. Fig 6 shows the average actin intensity profile along the channel at the point when the nucleus is entering into the constriction. For comparison the traction force magnitudes from the solid model (Fig 4b) and from shell model (Fig 5b) are plotted on the same graph. Of particular interest are the lateral forces around the middle of the nucleus corresponding to the peak in actin intensity around the beginning of the constriction. This strongly suggests that the increase in actin around the nucleus at the entrance to the constriction contributes to generating the force necessary to the deform the nucleus to pass it through the constriction. By this we therefore provide the first evidence that lateral actin exerts force deforming the nucleus from the sides. This lateral force is necessary for the nucleus to pass through such constrictions. Models such as [28] which assume axial forces only (either pushing or pulling along the axis) predict a threshold constriction size below which the nucleus would not be able to pass through. With the lateral forces our model predicts however the nucleus is able to pass through smaller constrictions. In addition the magnitudes of the forces we predict are lower than that required for a model with axial forces only. Thiam et al [17] show that lamin A/C depleted cells are able to pass through constrictions without lateral actin. Lamin A/C depletion softens the nucleus by disrupting the lamina and therefore the elastic shell of the nucleus. Without its elastic shell we expect the nucleus can pass through the constrictions with only axial forces because it is behaving more like a fluid.

**Fig 6.**
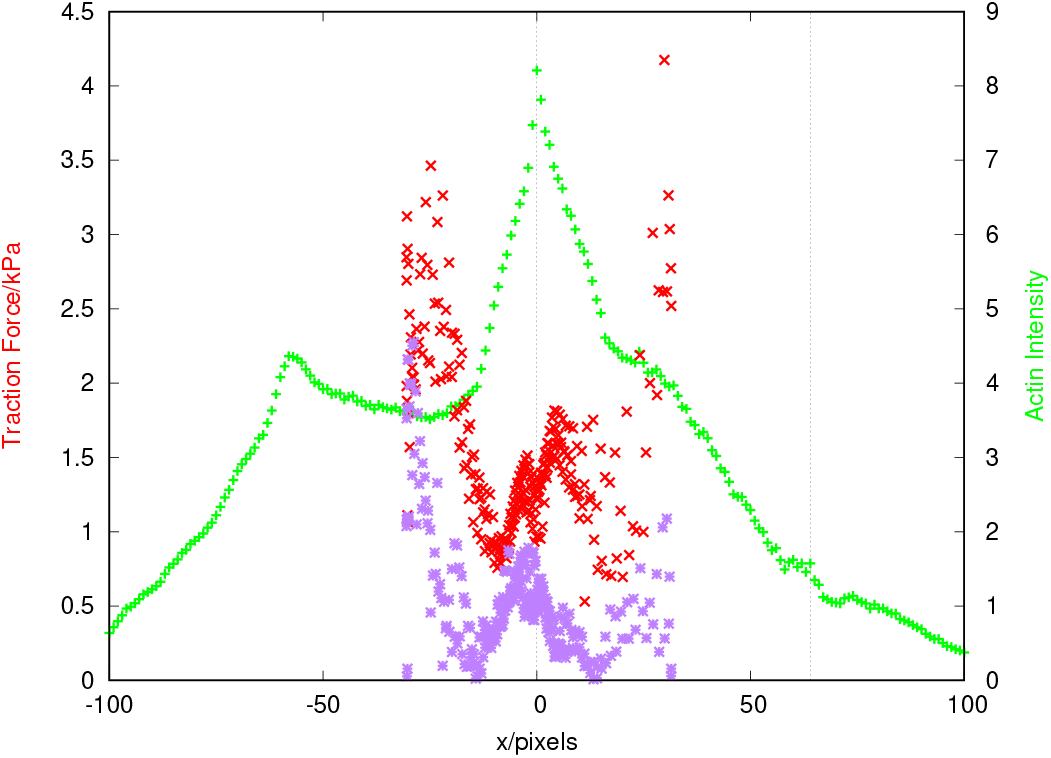
Average actin (LifeAct-GFP fluorescence) intensity (green + points) at the time point when the nucleus is entering into the constriction (right hand *y*-axis). The actin intensity is the mean intensity over the width of the channel at each pixel position. This is then renormalised by the average intensity for each cell and aligned with the start of the constriction at *x* = 0. The average is taken over 83 cells. The red × points are the traction force magnitudes from the solid model (Fig 4b) and the purple * points are the traction force magnitudes from shell model (Fig 5b).The vertical dotted line indicate the start and end positions of the constriction.

The two limiting cases of models we have studied here both show results indicative of experimental observations. In particular, the normal lateral forces predicted by the solid model correlate with actin density and RICM data from Thiam et al. [17] and the large extensile forces at the leading tip predicted by the shell model correlate with the tip rupture observed by Thiam et al., Denais et al. and Raab et al. [15–17]. We therefore expect that aspects of both models are important in nuclear deformation and that the actual viscoelastic properties of the nucleus lie between the two simple limiting elastic models we have presented here. The method and algorithm we present here for analysing images of nuclear deformation can be easily used with any chosen input model for the nucleus.

## 6 Conclusions

We have presented a method for calculating the force acting over the surface of a cell nucleus causing the deformation seen between two images, using a continuum elasticity model. We have used a Monte Carlo simulated annealing simulation method to define the deformation field between two images. We then calculated the traction force causing the observed deformation of averages of the nuclei of cells migrating through a constriction in a confined channel. We have presented two limiting case models for the nucleus as a homogeneous elastic solid and as a thin elastic shell. Since the most realistic model is likely to lie between these extremes, we expect the distribution of forces to be between the two cases we have calculated.

The predictions we have made of the details of the force fields required to deform a nucleus through a constriction can be used to assess models of force generation mechanisms used by cells to force their nucleus through small constrictions. For example the rearmost nuclear forces shown in figure 4(b-d) and 5(d) are consistent with what would be expected from actomyosin contraction at the rear of the cell generating an increased pressure behind the nucleus. The forces at the front of the nucleus may also be generated by such an actomyosin contractility generated pressure gradient or by pulling forces at the front of the cell, for example force exerted on the nuclear surface by molecular motors as they move along cytoskeletal filaments. By using our model, we were able to determine that the compressive forces perpendicular to the direction of motion seen in figure 4 correlate with the Arp2/3 mediated actin assembly seen by Thiam et al. [17] at the constriction entrance. We therefore conclude that actin polymerisation generates lateral forces around the nucleus that are required to squeeze it into constrictions that are smaller than the threshold size passable with axial forces alone. Our algorithm can be used to quantitatively analyse future experiments to further investigate such mechanisms used to generate nuclear deformation.

Our computational model is not restricted to nuclei within channels and could be applied to any two dimensional images of deforming nuclei, if appropriate assumptions are made. Furthermore our algorithm could be adapted to analyse images of other elastic objects deforming. Our code can be obtained freely on GitLab https://gitlab.com/nucleus-deformation-traction.

## Supporting information: Appendix

### 6.1 DNA density

Another method to determine whether there is any volume change of the nucleus within the channels is to observe the density of DNA within the cell. As DNA is the most dense component within the nucleus, observing how the density of the DNA changes with position in the channel could indicate if there are any significant volume changes. If the nucleus does increase in volume, the average density of materials within the nucleus, including the DNA should decrease and vice versa if the nucleus is compressed. Both density changes and height changes in the out of plane direction should change the observed florescence intensity of the DNA. Therefore, if we observe a change in fluorescence intensity by more than the change in the dimensions of the channel in to the constriction, that would be an indication of overall nuclear volume change.

In order to measure the average intensity of fluorescing pixels, we need to remove the background of the images. We did this using the existing software within imageJ [50], which uses a “rolling ball” method, in order to remove the background of images [56]. However, as in the example shown in [56], this typically leaves the background as pixels with small, but non zero values of fluorescence. In order to accurately count the number of fluorescing pixels, these must not be counted when computing the mean intensity value. To do this, we calculate the average intensity of all the pixels in a given image. We then calculate the mean value and standard deviation of the fluorescence of the pixels. Then we decrease the intensity of each image by an amount given by *I*_*new*_ = *I*_*old*_ − *Ĩ* − *aσ*, where *I*_*new*_ is the intensity with the background removed, *I*_*old*_ is the intensity with background, *Ĩ* is the mean intensity of all pixels, *σ* is the standard deviation of the intensities and *a* is a value that is dependent on the image. The value of *a* is increased in increments of 0.1 from 0, until the number of fluorescing pixels matches that of the area of the curves around the nuclei, which we had previously drawn, using the threshold tool and selection tools within imageJ, for each given image and frame.

The change in DNA intensity at each *x* coordinate as shown in figure 7 to between 60 − 70% is also consistent with the change in height of the channel from 5*μ*m outside of the constriction to 3.4*μ*m within the constriction, i.e. the constriction is approximately 70% of the height of the channel. We therefore conclude that there is no significant volume change of the nuclei as they travel through the constrictions.

**Fig 7.**
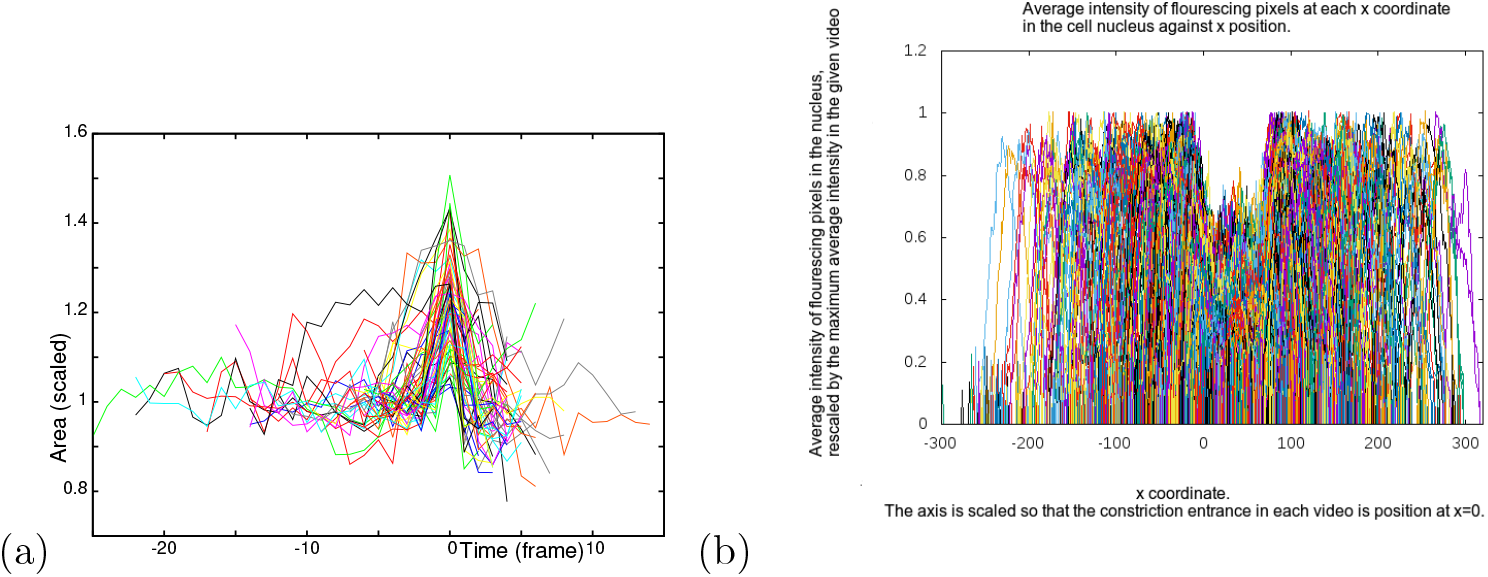
(a) The relative area change of dendritic cell nuclei as they travel through a channel of cross sectional shape 7*μm* by 5*μm* and a constriction of size 2*μm* by 3.4*μm*. The area of each nucleus is rescaled by the average area of the cell nucleus prior to entering the constriction. The time axis is centred on the peak value of area. The area increases as each nucleus passes through the constriction. (b) The average intensity of fluorescing pixels against position of 80 cell nuclei travelling through channels containing constrictions. The constriction entrance in each video is positioned at *x* = 0. The intensity of each video of a nucleus is renormalised by the maximum value of average intensity seen in the video.

### 6.2 Strain tensor in polar coordinates

In order to accommodate the solid model assumption of a deformation field decreasing lineally along the radial direction, the calculations of derivatives in this model are performed using polar coordinates in the 2d plane seen in images. However, the coordinates and deformation fields read in and calculated are known in Cartesian coordinates.

The polar form of the strain and stress equations can be written using

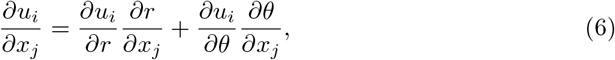

where we can use the standard Polar-Cartesian relations to evaluate the following terms;

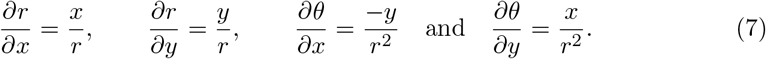

Substituting these into Eq (6) gives the polar form of the stress/strain tensor.

Numerically, we calculate these derivatives as given below, using the forms of numerical derivatives for varying spatial positions between mesh points.

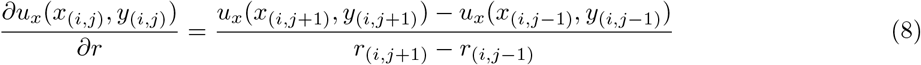

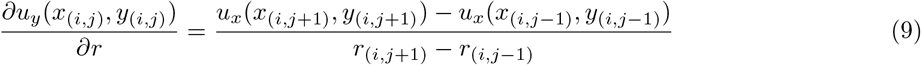

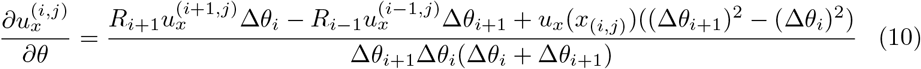

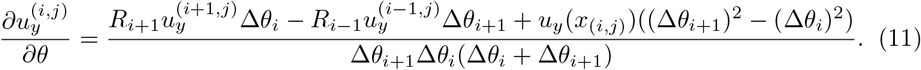

Where *R*_*i*+1_ is the ratio of the radius at point *i* + 1 to the radius at point *i*. Similar expressions are used for the innermost and outermost shapes, but replaced with forward/backward finite difference methods respectively. The factors of *R*_*i*_ in the final two equations are included to scale for small variations in the radius between points. These equations, together with the standard relations between Cartesian and polar coordinates allow the strains to be numerically evaluated from Eq (2) in the Cartesian coordinate basis directly.

To calculate derivatives of the deformation field in the solid model we need to define the deformation at points along a ring just inside the perimeter. Consider point *i* on the perimeter with coordinates (*x*[*i*]*, y*[*i*]) and position vector from the origin to the point *i* defined by ***r***[*i*] = (*x*[*i*]*, y*[*i*]). The corresponding point on the inner ring is defined as the point where ***r***[*i*] crosses the ring. The deformation at point *i* is (*u*_*x*_[*i*], *u*_*y*_[*i*]). We assume linearly elastic behaviour and zero deformation at the origin. This leads to the assumption that the deformation at the corresponding point on the inner ring is in the same direction as that at point *i* but with a linearly reduced magnitude. i.e. the deformation at the corresponding point on the inner ring is given by:

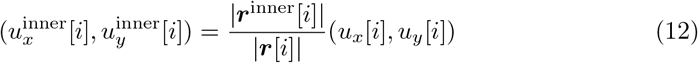

where |***r***^inner^[*i*]| is the distance from the origin to the inner ring point corresponding to point *i*. To calculate the derivatives of these deformations numerically we use a backwards difference method.

## 7 Differential geometry of surfaces

In order to generally describe the more complicated unknown surface in the general tangent and normal coordinate basis, and calculate values along the surface, we first describe the general form of the derivatives using differential geometry. The general forms involve the curvatures of the surface and the Christoffel symbols of the surface, and an analytic method to calculate the derivatives is given below.

Briefly, a surface ***X***(*n, s*_1_, *s*_2_) described by two tangential directions *s*_1_, *s*_2_ and the normal direction *n* has an associated metric tensor given by the derivatives of the surface along each of the directions at each point.

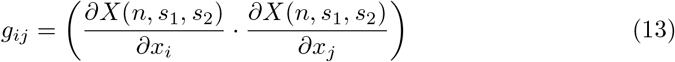

For a positively oriented surface, where by definition when travelling along the curve describing the surface, the interior of the curve is on the left, the outwards normal to the surface is then given by

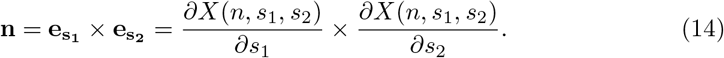

Then, for a surface with two tangent vectors **e** and **e′** with components in the basis (*x, y,* · · ·) represented by subscripts, the metric tensor is given as

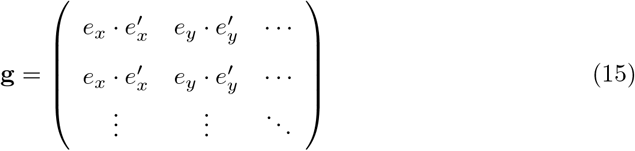

The Christoffel symbols are written in terms of the metric tensor as

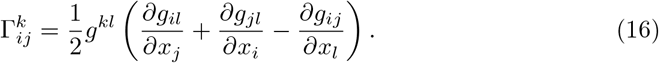

Similarly, the curvature of the surface, measured as the rate of change of the normal direction along the surface can be expressed as a tensor, *C*_*ij*_.

A thin shell surface can be written as a function of only the two tangent directions, ***X***(*n, s*_1_*, s*_2_), and so the metric tensor is a 2×2 matrix, with the components *s*_1_ and *s*_2_ representing two tangent directions along the surface.

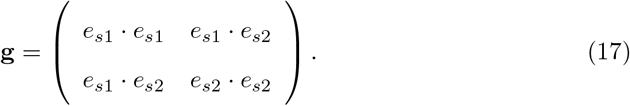

The derivatives along the surface of the basis vectors are then given in terms of the curvature and metric as

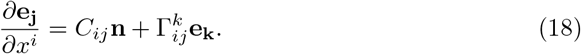

Using these relations, we can describe any thin shell surface, for instance those seen in images of the nucleus. Because the images considered are only in two dimensions, an analytical approach is used to describe the out of plane direction *s*_2_, while the in plane images provide *s*_1_ from the outline. The normal direction is assumed outwards and in the XY plane seen in images, and so can be determined purely from the *s*_1_ tangent vector. As the surface is flat in the out of plane direction, the vector in the out of plane direction is easily defined as a unit length vector parallel to the *z* axis. As such the metric tensor is the identity and the Christoffel symbols are all zero, leaving only the curvature terms in the shell model of this particular out of plane direction shape. However, we include the full differential geometry in the code to allow for use with other, more complicated, shapes beyond the scope of this paper.

## 8 Alignment of initial and target images

The deformation free point is likely to lie somewhere between the two limiting cases described in section 4.2. However the limits on both the spatial and temporal resolution of the nuclei in the images prevent the point of zero deformation from being identified directly from any given series of images.

In order to estimate the location where the deformed nucleus should be placed relative to the undeformed nucleus, we measured how the position of the front and rear of the nucleus changed between frames. Measurements of the change in the front and rear position of each nucleus were taken between frames at each of the five positions (before the constriction, entering in to the constriction, inside the constriction, leaving the constriction and after the constriction).

Fig 8 shows the changes in position of the rear of each nucleus, against the change in position of the front of the same nucleus, with a linear fit of the form *y* = *mx* + *c*. The gradient of the lines of best fit, as shown in each of the graphs, provides an estimate of the how far the front position of the nucleus will move, given a change in the rear position, or vice versa. The intercept with the *y* axis measures how much the rear of the nucleus will move when the front of the nucleus does not change position. The *y* intercept is near zero in figures 8a and 8b, consistent with the nucleus being unable to move the rear without the front of the nucleus deforming as it entering the constriction, as the out of plane direction is already filled by the nuclei volume. In figures 8c and 8d, the *y* intercept is larger than from zero, representing the nucleus filling the volume in the out of plane direction and unlike the entry position the nucleus can move freely into the larger space post-constriction.

**Fig 8.**
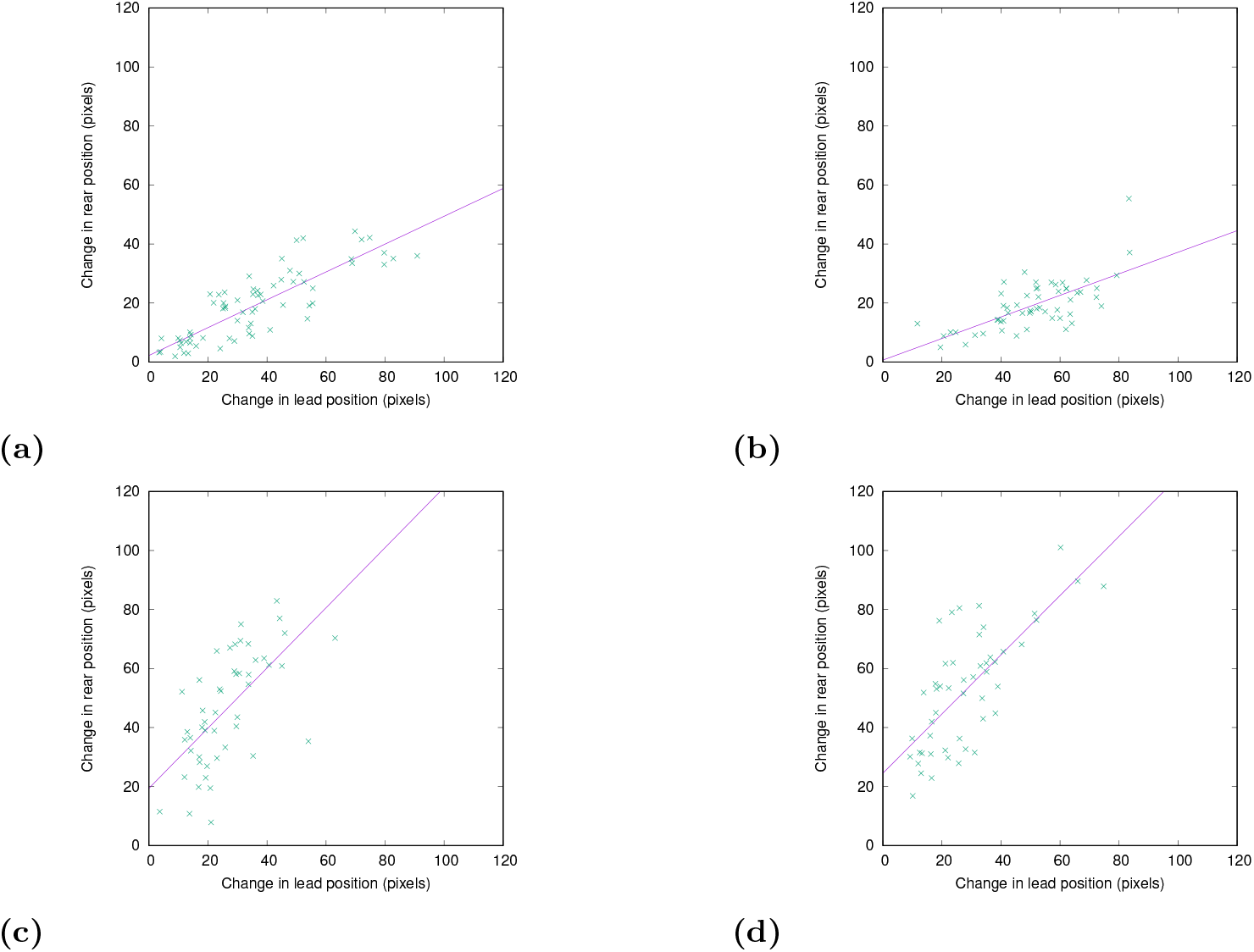
These figures show the change in the rear position of each nucleus against the change in leading position of the same nucleus. Each point in each of the graphs represents one nucleus. (a) shows the changes as the nucleus moves from before the constriction to beginning to enter the constriction. (b) shows the changes from when the nucleus is entering the constriction, to when it is fully in the constriction. (c) shows when the nuclei are moving from in the constriction to leaving the constriction. Finally, (d) shows the nuclei as they go from leaving the constriction to having fully exited the constriction. The best fit lines in each image are for (a): *y* = 0.47(±0.04)*x* + (2.23 ± 1.48), for (b): *y* = 0.37(±0.05)*x* + 0.68(±2.81), for (c): *y* = 1.02(±0.18)*x* + 19.4014(±5.179) and for (d): *y* = 1.00(±0.1375)*x* + 24.52 ± 4.38.

The gradient provides an estimate of where the point of zero deformation should be between each average deformation of the nucleus. The nuclei are initially aligned by the centres of mass, and then shifted an amount along the x axis, to reflect the change given by the ratio of the change in position of the rear to the change in position of the front of the nuclei. As the nuclei are orientated so that they all move in the positive *x* direction, the value of the ratio is always positive. If the ratio of the change in the rear position to the change in the front position, *m*, is *m* ≥ 1, then the rear moved more than the front, and the target nucleus shape is shifted forwards in space relative to the undeformed nucleus. Vice versa if the ratio is in the range 0 ≤ *m* ≤ 1, then the nucleus is shifted in the opposite direction.

The distance that we shift the entire target shape along *x* is proportional to the relative change in the rear or front position compared to the sum of the changes to the rear and the front in the respective directions. The proportion of the distance to move in the given direction is,

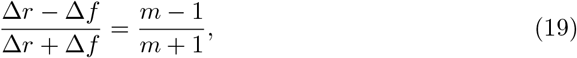

where Δ*r* and Δ*f* are the changes in the rear/front of the nucleus position between the undeformed and deformed shapes when aligned in the centre of mass frames respectively. The values used to shift each target shape are given in table 1.

**Table 1.** The changes in position used to shift the target shape. Δ*r* and Δ*f* are the changes between the rear and front position of the nuclei respectively. All distances are in pixels.

| Position | Gradient | Shift of nucleus position<br>from centre of mass |
| --- | --- | --- |
| Before to entering | 0.47 | $-0.36\vert \Delta r\vert = 1.7$ |
| Entering to in | 0.37 | $-0.46\vert \Delta r\vert = 7.3$ |
| In to exiting | 1.02 | $0.01\vert \Delta f\vert = 0.15$ |
| Exiting to out | 1.00 | $0.00\vert \Delta f\vert = 0.0$ |

## Acknowledgements

We acknowledge the EPSRC Standard Research Studentship DTG for IDE, EPSRC grant no. EP/L026848/1 and the University of Sheffield.

